# A *Mycobacterium tuberculosis* effector targets mitochondrion, controls energy metabolism and limits cytochrome c exit

**DOI:** 10.1101/2021.01.31.428746

**Authors:** Marianne Martin, Angelique deVisch, Yves-Marie Boudehen, Philippe Barthe, Claude Gutierrez, Obolbek Turapov, Talip Aydogan, Laurène Heriaud, Jerome Gracy, Olivier Neyrolles, Galina V. Mukamolova, François Letourneur, Martin Cohen-Gonsaud

## Abstract

Host metabolism reprogramming is a key feature of *Mycobacterium tuberculosis (Mtb)* infection that enables the survival of this pathogen within phagocytic cells and modulates the immune response facilitating the spread of the tuberculosis disease. Here, we demonstrate that a previously uncharacterized secreted protein from *Mtb,* Rv1813c manipulates the host metabolism by targeting mitochondria. When expressed in eukaryotic cells, the protein is delivered to the mitochondrial intermembrane space and promotes the enhancement of host ATP production by boosting the oxidative phosphorylation metabolic pathway. Furthermore, the release of cytochrome c from mitochondria, an early apoptotic event in response to short-term oxidative stress, is delayed in Rv1813c expressing cells. This study reveals a novel class of mitochondria targeting effectors from *Mtb* which might participate in host cells metabolic reprogramming and apoptosis control during *Mtb* infections.

## Introduction

*Mycobacterium tuberculosis (Mtb)* encodes secreted virulence factors contributing to its successful infection of host cells and its ability to actively replicate inside the phagosome (Hmama et al., 2015) (Winden et al., 2019). After phagocytosis, *Mtb* blocks phagosomal maturation, escapes phagosomes and subverts the host immune response. Several virulence factors *(e.g.* proteins, lipids) have been already described to mediate such mechanisms, but corruption of host cell defenses is clearly multifactorial (Nicholson et al., 2021). It is estimated that over 20% of bacterial proteins have functions outside the bacterial cytoplasm and are exported to their designated locations by protein export systems (Kostakioti et al., 2005). Identification of secreted proteins remains a challenging task (Målen et al., 2007; de Souza et al., 2011; Albrethsen et al., 2013). The comparison of *Mtb* secreted proteins reported in various proteomic studies revealed only a small number of proteins consistently identified (Tucci et al., 2020). As experiments were made in various culture conditions, it is not surprising that secretion patterns differ from one experiment to another. Furthermore, the host cell environment also plays an important role in defining the secretion pattern, as recently revealed by studies focusing on the identification of secreted proteins during infection (Perkowski et al., 2017; Penn et al., 2018). To get a broader view on the *Mtb* secretome, we used multidisciplinary approaches including bioinformatics, structural and biochemical techniques, and cellular biology analyzes. We identified putative *Mtb* secreted proteins using proteins primary sequence analysis combined with structure modelling. Among the selected targets, we studied the protein coded by the *rv1813c* gene which is only present in mycobacterial pathogens. The Rv1813c protein has been used as vaccine adjuvant (Bertholet et al., 2008) and displays immunogenicity properties (Liang et al., 2019). Rv1813c expression was reported to be MprA and DosR regulated (Bretl et al., 2012) and *Mtb* ΔRv1813c mutant was attenuated in the low-dose aerosol model of murine tuberculosis (Bretl et al., 2012).

In this paper, we describe molecular and functional analyzes of this protein. We showed that Rv1813c defines a new class of effectors, with an original fold, addressed to mitochondria. Mitochondrion plays critical functions not only supplying cells with energy but also contributing to several cellular mechanisms including cell cycle, apoptosis, and signaling pathways. Metabolism modulation dictates macrophage function and subsequent *Mtb* infection progression. Here, we demonstrate that Rv1813c affects some mitochondrial metabolic functions and cellular responses to oxidative stress. These results suggest that Rv1813c might play regulatory roles in the metabolic and apoptotic responses occurring in *Mtb* infected macrophages.

## Results

### Bioinformatic analysis of *Mtb* genome for identification of secreted proteins

*Mtb* possesses at least three different secretion systems using distinct secretion determinants (structural and/or motif-based) present on transported proteins (Feltcher et al., 2010). To predict secreted proteins *in silico,* we analyzed the predicted *Mtb* H37Rv proteome using an in-house Protein Analysis Toolkit (PAT) (Gracy and Chiche, 2005). First, SignalP v4.1 and PredTAT softwares were used to predict the presence of known signal peptides and/or structural features necessary for secretion. In addition, transmembrane segments were inferred using either Uniprot annotations or the TmHMM prediction software. The number of predicted transmembrane segments and the position of the last transmembrane segment were also manually examined to identify signals potentially missed by the other methods. To search for potential T7SS-mediated secreted proteins, we first performed helix structure prediction of each protein using Psipred (McGuffin et al., 2000), and then searched for the YxxxD/E motif in between the characteristic two helices (Daleke et al., 2012). These data were compared with various proteomic data and model databases (ModBase, Interpro, GO). Among the proteins identified here as potentially secreted, we studied Rv1813c, a 143 amino-acid protein with a predicted folded domain of unknown function.

### Rv1813c protein sequence features and secretion

Primary sequence analysis of the Rv1813c protein unambiguously identified a potential signal sequence (residues 1 to 28) with an upstream arginine repeat (residues 6-8) indicating that the protein could be exported by the Tat export system (**Fig. 1A**). Consistent with its genuine export signal, Rv1813c has been identified in one culture filtrate proteome (Tucci et al., 2020), but surprisingly is absent in three others published secretomes (Malen et al., 2007; de Souza et al., 2011; Albrethsen et al., 2013). Conversely, Western blot analysis of *Mtb* culture filtrates using a specific antibody here confirmed that Rv1813c is secreted during active growth in culture medium (**Supplementary Fig. S1**). Rv1813c homologous proteins are mostly found in Actinobacteria *(Mycobacterium, Nocardia* and *Streptomyces* genera). In addition to *Mtb,* the protein is present in various mycobacteria including *Mycobacterium marinum (Mmar), Mycobacterium avium, Mycobacterium ulcerans* and *Mycobacterium abscessus.* Multiple paralogues exist within the same bacteria. For instance, *Mtb* possesses only one orthologue (Rv1269c), whereas *Mmar* harbors three paralogues (MMAR_1426, MMAR_2533 and MMAR_4153). Remarkably, Rv1269c has been detected in all culture filtrates proteomes published so far (Malen et al., 2007; de Souza et al., 2011; Albrethsen et al., 2013). Therefore, the secretion in culture medium might be a common feature of Rv1813c homologous proteins. The sequence similarity between these various proteins is high (between 45 to 70%), with a lower sequence identity for the N-terminal part (residues 28-54 for Rv1813c) of the protein (**Fig. 1A**). Four cysteine residues are present and conserved. The last four amino acid residues (^140^WACN^143^) composed a strictly conserved motif in all Rv1813c homologous proteins that includes one of the conserved cysteines. Fold-recognition and modelling server @TOME2 previously used in many studies for protein function identification even at low sequence identity (Turapov et al., 2014) failed to identify any close or distant Rv1813c structural homologues.

**Fig. 1:**
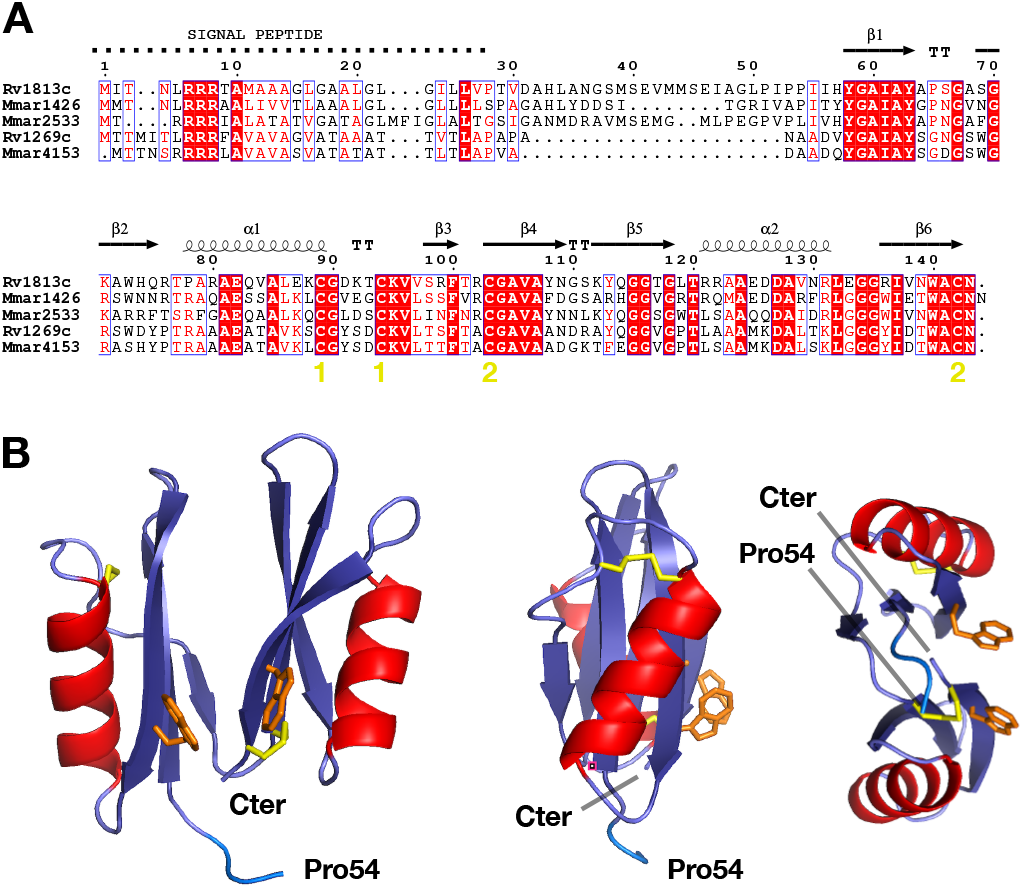
Rv1813c defines a new protein family. **(A)** *M. tuberculosis* and *M. marinum* Rv1813c homologues sequence alignments. The secondary structure of Rv1813c is reported above the alignment. The yellow numbers indicate the cysteine engaged in disulfide bridges. **(B)** Rv1813c structure determined by multi-dimensional NMR. Three cartoon representations of the structure. Only four residues of the N-terminal unfolded part of the protein (residues 28-57) are represented. The cysteine residues, all engaged in disulfide bridges are represented in yellow while the two solvent-exposed tryptophan amino acids are represented in orange.

### Rv1813c defines a new protein family and a unique protein fold

The Rv1813c-coding sequence without the first codons corresponding to the protein signal peptide (residues 1 to 27) was cloned into an *Escherichia coli* expression vector. The protein was over-expressed as inclusion bodies, purified and refolded. The purified protein was used for multidimensional NMR experiments. Preliminary examination of [1H,15N]-HSQC spectrum revealed that 30 residues were unfolded (**Supplementary Fig. S2**). A full multi-dimensional NMR study led to the protein three-dimensional structure resolution (**Fig. 1B, Supplementary Table S1**). Structure resolution demonstrated that the residues 28 to 57 were unfolded and that the protein possessed a 86 residues folded domain at its carboxyl terminus. This domain is composed of two duplicate lobes facing each other, and each lobe comprises a series of three *ß*-strands with a hydrophobic surface and an *α*-helix (*ß*/*ß*/*α*/*ß*). The four conserved cysteines are engaged in two disulfide bonds located in different parts of each lobe. The proteins from the family defined by the Rv1813c sequence (Pfam domain: DUF4189) contain a conserved WACN motif (Trp-Ala-Cys-Asn residues) with the cysteine engaged in a disulfide bond linking the strands *ß6* and *ß*4, while its tryptophan is solvent-exposed as well as the second tryptophan (Trp140) in Rv1813c. In addition to protein folding and stability properties, these solvent-accessible tryptophan residues might be functionally important for Rv1813c, contributing either to an hypothetic Rv1813c active site or to interactions with other ligands as classically described for solvent-exposed residues. The overall structure defines a previously undescribed fold as both Dali (Holm and Rosenström, 2010) and FATCAT (Ye and Godzik, 2004) servers failed to detect any structural homologues. Consequently, sequence and structure comparison analysis did not bring any indication on the potential biological function of the Rv1813c protein family.

### Rv1813c is addressed to mitochondria in *Dictyostelium* cells

To assess the function of Rv1813c in host cells, we first used the amoeba *Dictyostelium dis-coideum.* This professional phagocyte is amenable to biochemical, cell biological and genetic approaches, and has proved to be an advantageous host cell model to analyze the virulence of several pathogenic bacteria (Steinert, 2011; Müller-Taubenberger et al., 2013). Furthermore, the intracellular replication of *Mmar* has been extensively studied in *D. discoideum* and shows similarity to *Mtb* replication in macrophages (Cardenal-Muñoz et al., 2017), indicating that comparable molecular mechanisms are at play in infected *D. discoideum* and mammalian cells. We first analyzed the intracellular localization of Rv1813c when over-expressed in *D. discoideum* (ectopic expression). Though protein expression levels might differ from what is encountered during *Mtb* infection, ectopic expression allows the advantageous analysis of individual secreted mycobacterial proteins without the potentially complexity brought by other bacteria effectors. Hence, this unphysiological expression of Rv1813c serves as a first step towards the biological characterization of this protein and might give some hints on its potential function(s) during *Mtb* infections.

Rv1813c deleted of its predicted signal peptide (first 27 amino acid residues) was tagged with a myc epitope at its N-terminus (myc-Rv1813c*_*P28-N143, here after referred to as myc-Rv1813c) and stably expressed in *D. discoideum.* Confocal microscopy analysis revealed colocalization in ring like structures of myc-Rv1813*c* coinciding with a mitochondrial outer membrane protein, Mitoporin (Troll et al., 1992) **(Fig. 2A).** Mitochondrial targeting was also observed in cells expressing Rv1813c tagged at the C-terminus (Rv1813c-myc) but was lost when Rv1813c was fused to GFP (**Supplementary Fig. S3A**). This specific targeting was independent of the added myc-tag as staining with an anti-Rv1813c polyclonal antibody of untagged Rv1813c showed identical results (**Supplementary Fig. S3B**). Mitoporin staining patterns were similarly observed in both parental (Ax2) and Rv1813c transfected cells (**Supplementary Fig. S3B, S3C**) excluding gross mitochondrial morphological defects induced by Rv1813c expression in *D. discoideum.* In cells labeled with mito-tracker deep red, a specific dye accumulating inside mitochondria, myc-Rv1813c surrounded labeled mitochondria and was mostly excluded from internal structures (**Fig. 2B**). This result suggested that Rv1813c might be attached either to the internal or the cytosolic sides of mitochondrial outer membranes. Interestingly, deletion of the unfolded N-terminus region of Rv1813c (deletion of residues 28 to 48; myc-Rv1813c_49-143) had no effect on Rv1813c localization whereas Rv1813c deprived of the folded region (deletion of residues 57 to 143; myc-Rv1813c_28-56) was not transported to mitochondria (**Fig. 2C**). Thus, the Rv1813c folded domain, which does not contain any known mitochondrial targeting signals, was sufficient to specifically direct this protein to mitochondrial outer membranes, whereas the unfolded N-terminus region appeared to be dispensable for Rv1813c targeting to mitochondria.

**Fig. 2:**
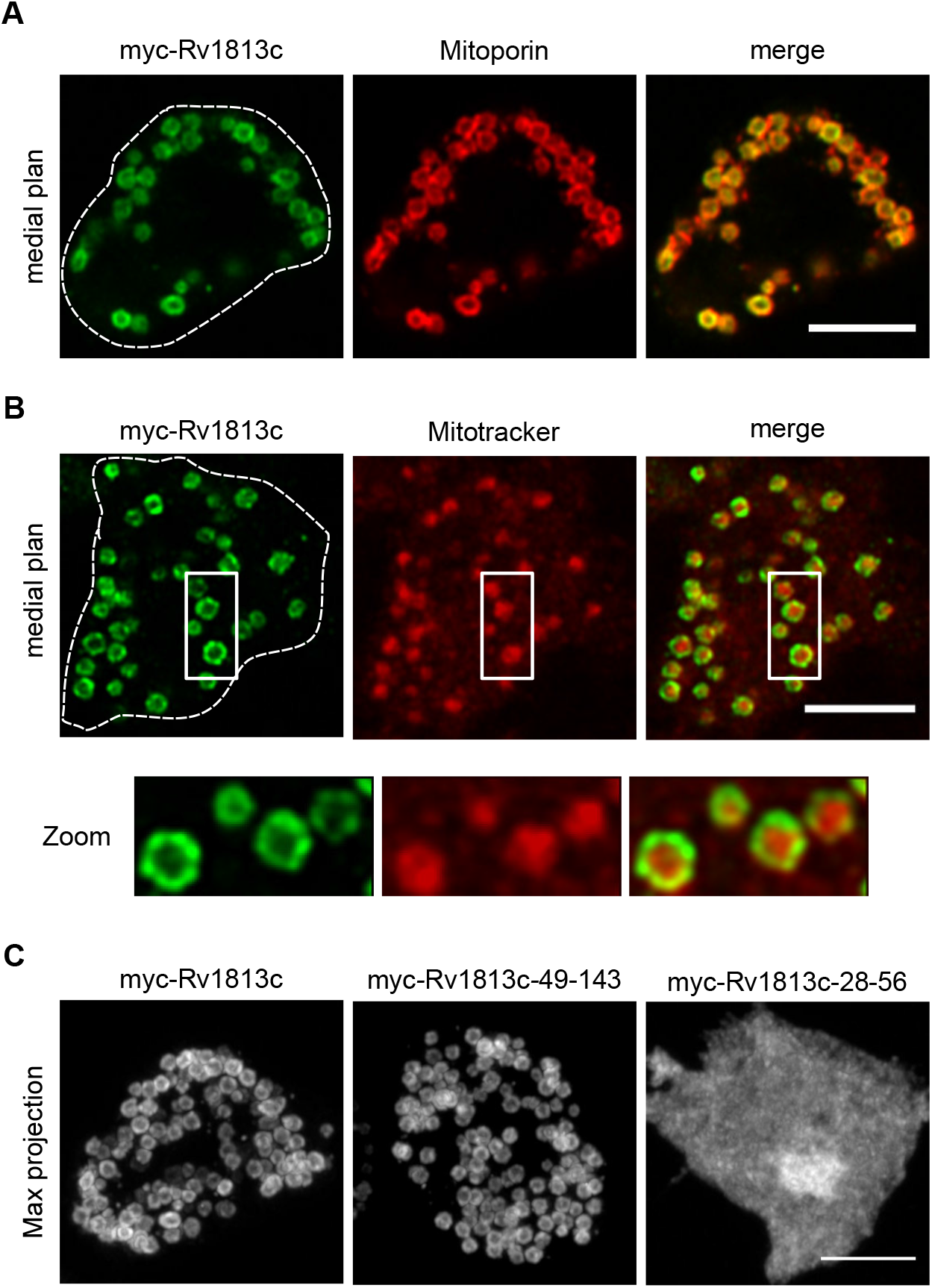
Mitochondrial localization of Rv1813c in *Dictyostelium.* *Dictyostelium* cells expressing the indicated constructs were fixed, processed for immunofluorescence, and analyzed by confocal microscopy (Airyscan). **(A)** Colocalization of myc-Rv1813c (detected with a rabbit polyclonal to Rv1813c) and mitochondrial Mitoporin in ring shaped structures. **(B)** Labeling of mitochondria of the indicated cells lines with Mitotracker deep red dye. Close-ups of mitochondria are shown in the inserts. **(C)** Maximum fluorescence intensity projection of Z confocal sections of cells expressing the full-length protein (myc-Rv1813c), the sole structured domain (myc-Rv1813c_49-143) and the unfolded domain (myc-Rv1813c_28-56) labeled with an anti-myc antibody. Cell contours are indicated by dotted lines. Bar, 5 μm.

### Rv1813c homologues are addressed to mitochondria in *Dictyostelium* cells

Intracellular localization studies were next extended to members of the Rv1813c family in *Mtb* and *Mmar* in the amoeba. All these proteins were detected in mitochondria, however some Rv1813c-like proteins affected mitochondria morphology. Whereas the overexpression of Rv1813c *Mmar* orthologs (MMA_1426 and MMA_2533) did not induce any apparent morphological defects in mitochondria, cells expressing Rv1269c or its *Mmar* ortholog MMA_4153 displayed mitochondria with aberrant shapes and sizes (**Supplementary Fig. S3D**). In addition to mitochondria, MMA_4153 also localized to the cytosol. Taken together, these results indicated that mitochondrial targeting in *D. discoideum* is a characteristic feature of the Rv1813c family, and for some members, this localization leads to defective mitochondrial morphology.

### Rv1813c resides in the mitochondrial intermembrane space

Mitochondria are composed of two membranes, the outer and inner membranes, separated by an intermembrane space (IMS**) (Fig. 3A)**. To determine more precisely the localization of Rv1813c within these submitochondrial compartments in *D. discoideum*, we next applied a biochemical approach. First, mitochondria enriched fractions (here after referred to as mitochondria) were obtained by subcellular fractionation (see scheme **Fig. 3B**). As expected, Rv1813c was recovered from the mitochondrial fraction confirmed by Mitoporin enrichment (**Fig. 3C**). Next, Triton X-114 phase partitioning experiments revealed that Rv1813c is not an integral membrane protein, in agreement with the absence of any predicted transmembrane domains (**Fig. 3D**) and its exclusion from the *Mtb* cell wall **(Supplementary Fig. S1)**. Consistently, Rv1813c was extracted from mitochondrial membranes by sodium carbonate treatment, a characteristic feature of peripheral membrane proteins (**Fig. 3E**). Since Rv1813c was not released from mitochondria by high salt washes (**Fig. 3F**) and was protected from proteinase K digestion of intact mitochondria (**Fig. 3G**), we concluded that Rv1813c resides inside mitochondria. In addition, Rv1813c was partially released from mitochondria upon the specific rupture of mitochondrial outer membranes in hypotonic buffer indicating that Rv1813c accumulates into the mitochondrial IMS where it could be weakly attached to the internal or external sides of mitochondrial outer or inner membranes respectively (**Fig. 3H**).

**Fig. 3:**
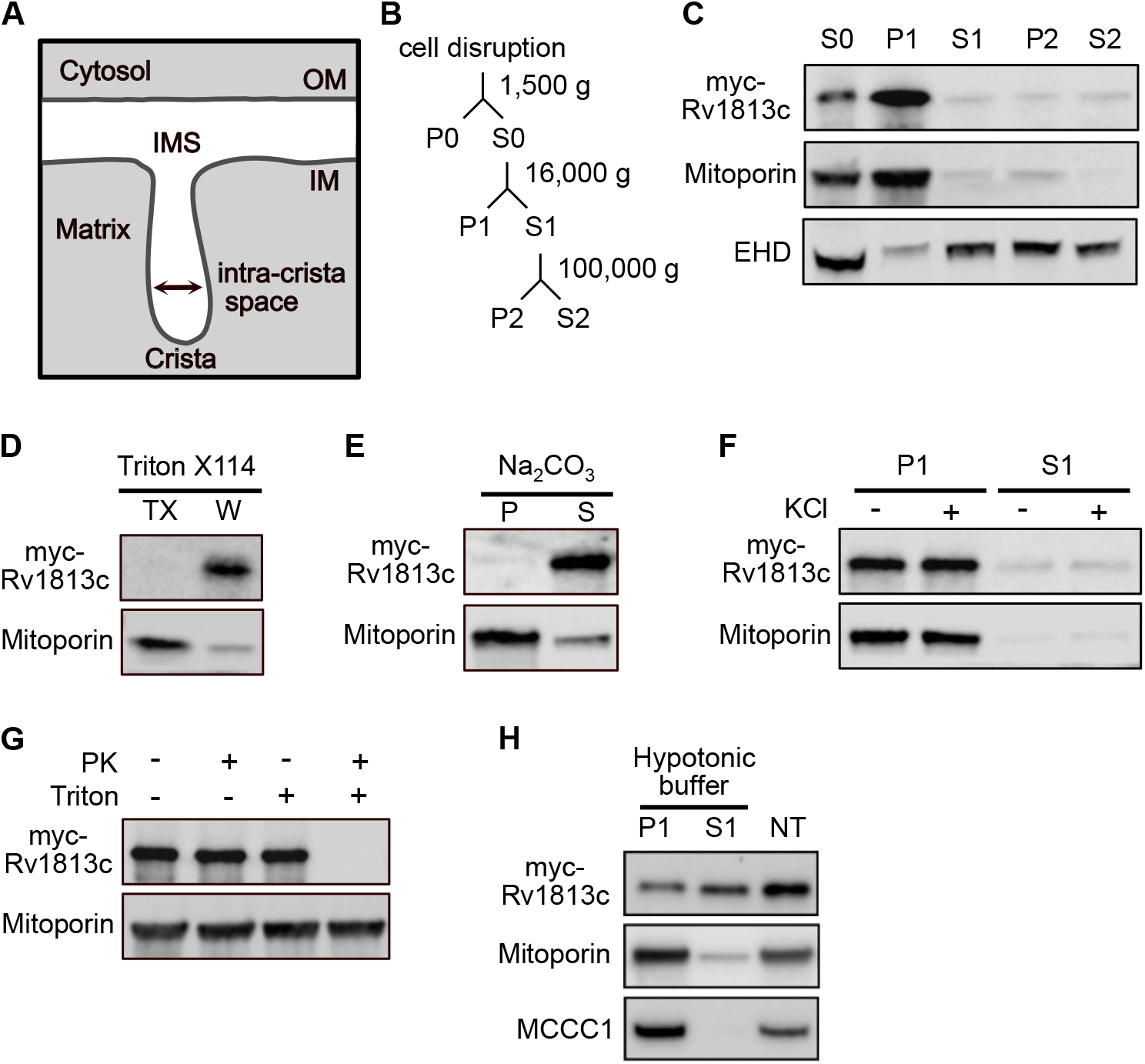
Biochemical analysis of Rv1813c mitochondrial localization in *Dictyostelium.* **(A)** Schematic ultrastructure of a single crista in mitochondria. Abbreviations: IMS, intermembrane space; IM, inner membrane; OM, outer membrane. (**B**) Fractionation scheme of differential centrifugation steps used to purify the Rv1813c enriched fraction from *Dictyostelium* cells. P and S represent pellet and supernatant, respectively **(C)** Fractions were analyzed by immunoblotting with antibodies to Mitoporin (mitochondria), EHD (endocytic vacuoles) and myc-tag (myc-Rv1813c). **(D)** The mitochondrial fraction was fractionated by Triton X114 extraction. The separated Triton X-114 (TX) and aqueous (W) phases were analyzed as above. Rv1813c is not extracted by Triton X-114 indicating no insertion inside membranes. **(E)** Mitochondria were incubated in sodium carbonate for 30 min. Rv1813c is mainly detected in the supernatant fraction (S) after centrifugation at 100,000 g of treated mitochondria, a characteristic of soluble and/or membrane peripheral proteins. **(F)** For high-salt protein extraction, mitochondria were incubated in buffer ± 200 mM KCl for 30 min and centrifuged at 16,000 g for 10 min. **(G)** Intact or Triton X100 treated mitochondria were subjected to proteinase K digestion for 30 min and analyzed by immunoblotting. **(H)** Mitochondria swelling was induced by hypotonic buffer incubation for 30 min. Released proteins (S1) from broken outer membranes (P1) were recovered by centrifugation at 16,000g for 10 min and analyzed by western blotting to detect myc-Rv1813c, Mitoporin (mitochondrial outer membrane) and mitochondrial 3-methylcrotonyl-CoA carboxylase α (MCCC1; mitochondrial matrix). Non-treated (NT) mitochondria incubated for 30 min in mitochondria isolation buffer A served as release specificity control.

### Rv1813c and orthologous proteins are addressed to mitochondria in mammalian cells

We next extended the analysis to mammalian cells. Native and myc-tagged Rv1813c were transiently expressed in HeLa cells and their intracellular localization was determined by confocal microscopy. As observed in *Dictyostelium,* Rv1813c was efficiently targeted to mitochondria in HeLa cells (**Fig. 4A and Supplementary Fig. S4**) without any detectable effects on the mitochondrial morphology (**Fig. 4B**). MMA_1436 and MMA_2533, two *Mmar* orthologues of Rv1813c also localized to mitochondria. However, in contrast to *Dictyostelium* cells, Rv1269c remained in the cytosol similarly to MMA_4153, the *Mmar* orthologs of Rv1269c, which also showed a faint mitochondrial localization (**Supplementary Fig. S4**). Although it might denote an intrinsic feature, we cannot rule out that the folding of these proteins might not proceed appropriately in mammalian cells impeding their efficient mitochondrial localization.

**Fig. 4:**
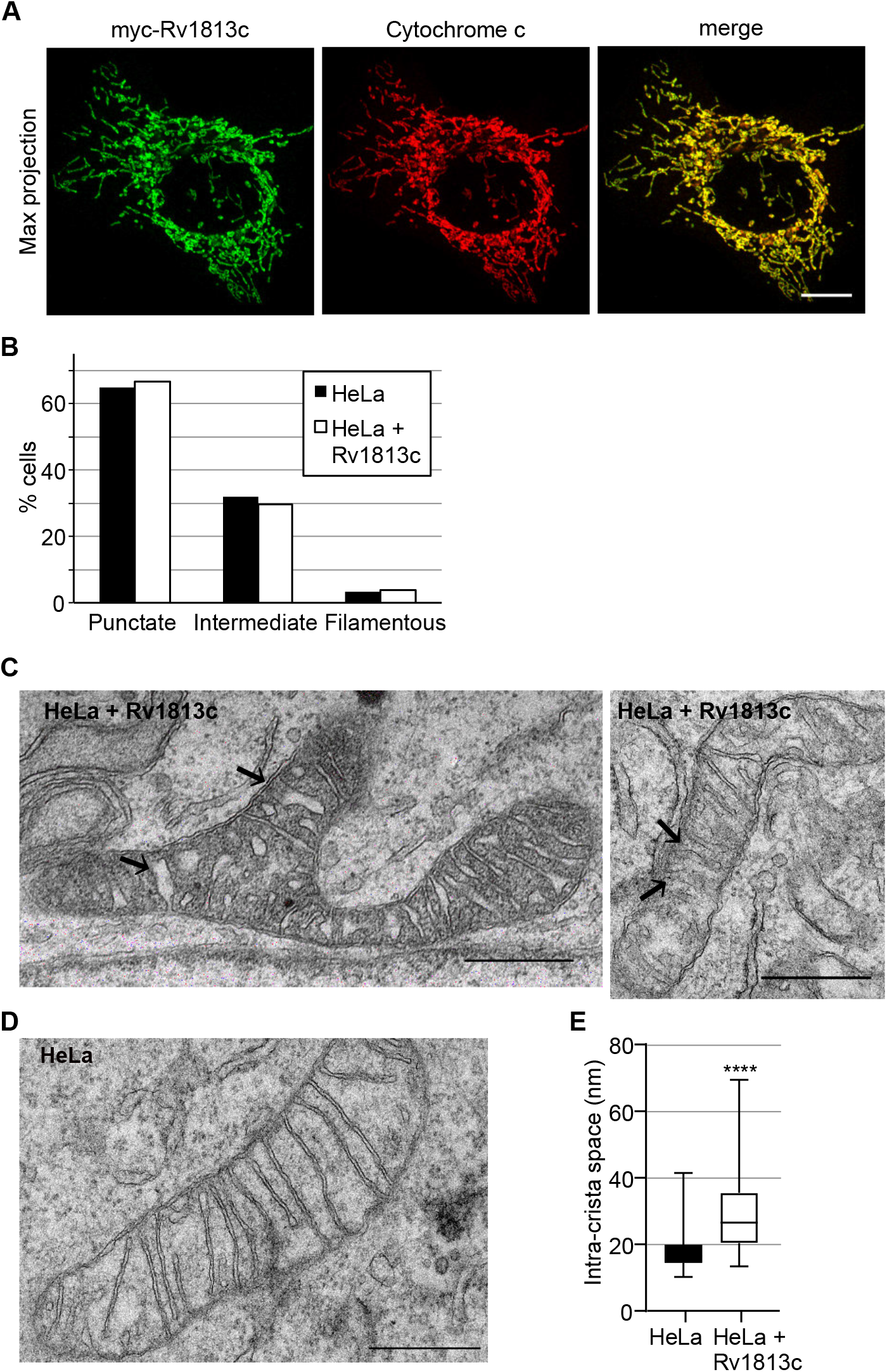
Targeting of Rv1813c to mitochondria in mammalian cells. **(A)** Confocal microscopy analysis of HeLa cells transiently expressing myc-Rv1813. Cells were fixed 48h post-transfection, processed for immunofluorescence with antibodies to RV1813c and cytochrome c, and analyzed by Airyscan microscopy. Bar, 10 μm. **(B)** Quantitative analysis of the mitochondria morphology observed in HeLa cells transiently transfected with pCI-myc-Rv1813 or empty vector as control. Mitochondria morphology was manually identified by confocal microscopy and classified in one hundred cells of one representative experiment from three independent analyzes. **(C, D)** Representative mitochondria ultrastructure determined by transmission electronic microscopy of MACS enriched HeLa cells transiently transfected with pMACS-4-IRES-II Rv1813c (C) or vector alone (D). Black arrows indicate some enlarged intra-cristae spaces in Rv1813c expressing cells. Bars, 500 nm. **(E)** Box-plot of intra-cristae spaces measurements for the indicated cell lines (100 random measurements each, **** p<0.0001 in student test). The bar inside the box indicates the median value. The bottom and top edges of the box indicate the 25th and 75th percentiles, respectively. The whiskers extend to the most extreme data points not considered outliers.

Whereas the overall morphology of mitochondria was preserved upon Rv1813c ectopic expression, transmission electronic microscopy (TEM) revealed some ultrastructural modifications. Hence, Rv1813c expressed in HeLa cells contained mitochondria with either normal or electron-dense matrix, and the intra-cristae space appeared significantly enlarged compared to parental HeLa cells **(Fig. 4C, 4D).** This altered mitochondrial morphology is reminiscent to what is observed in *Mtb* infected macrophages (Abarca-Rojano et al., 2003). Since cristae membranes are enriched in resident proteins involved in oxidative phosphorylation, this particular ultrastructure might lead to several mitochondrial energetic/metabolism consequences.

### Rv1813c overexpression enhances cell metabolism and mitochondrial ROS production

Next, we investigated whether changes in the mitochondrial morphology might cause energy metabolism disorders. Oxidative phosphorylation (OXPHOS) and glycolysis were simultaneously analyzed in intact cells making use of an extracellular flux analyzer (XF, Agilent Seahorse). In this assay, mitochondrial respiratory characteristics are evaluated by recording oxygen consumption rate (OCR) upon sequential chemical perturbation of selected mitochondrial functions (as detailed in Fig. 5 legend). In Rv1813c transfected HeLa cells, basal respiration, ATP-linked respiration, maximal respiratory capacity and reserve capacity were significantly increased compared to parental HeLa cells (**Fig. 5A-B**). Glycolysis was also assayed using a glycolysis stress test (Agilent Technologies) and measurements of extracellular acidification rates (ECAR) in incubation media. This assay revealed similar glycolytic profiles in control and Rv1813c expressing HeLa cells (**Fig. 5C-D**). Next, mitochondrial membrane potential was tested using flow cytometry of JC-1 stained cells. This assay revealed that expression of Rv1813c in Hela cells had no effect on ΔΨ_M_ in resting cells (**Fig. 5E**). However, these cells showed a slight but significant increased mitochondrial ROS production (**Fig. 5F**). Taken together, these results indicate that Rv1813c expression improves mitochondrial respiratory capacities without altering glycolytic functions, driving cells into an energy activated state. This higher mitochondrial respiration is associated with a moderate increased mitochondrial free radical formation without changes in the mitochondrial membrane potential.

**Fig. 5:**
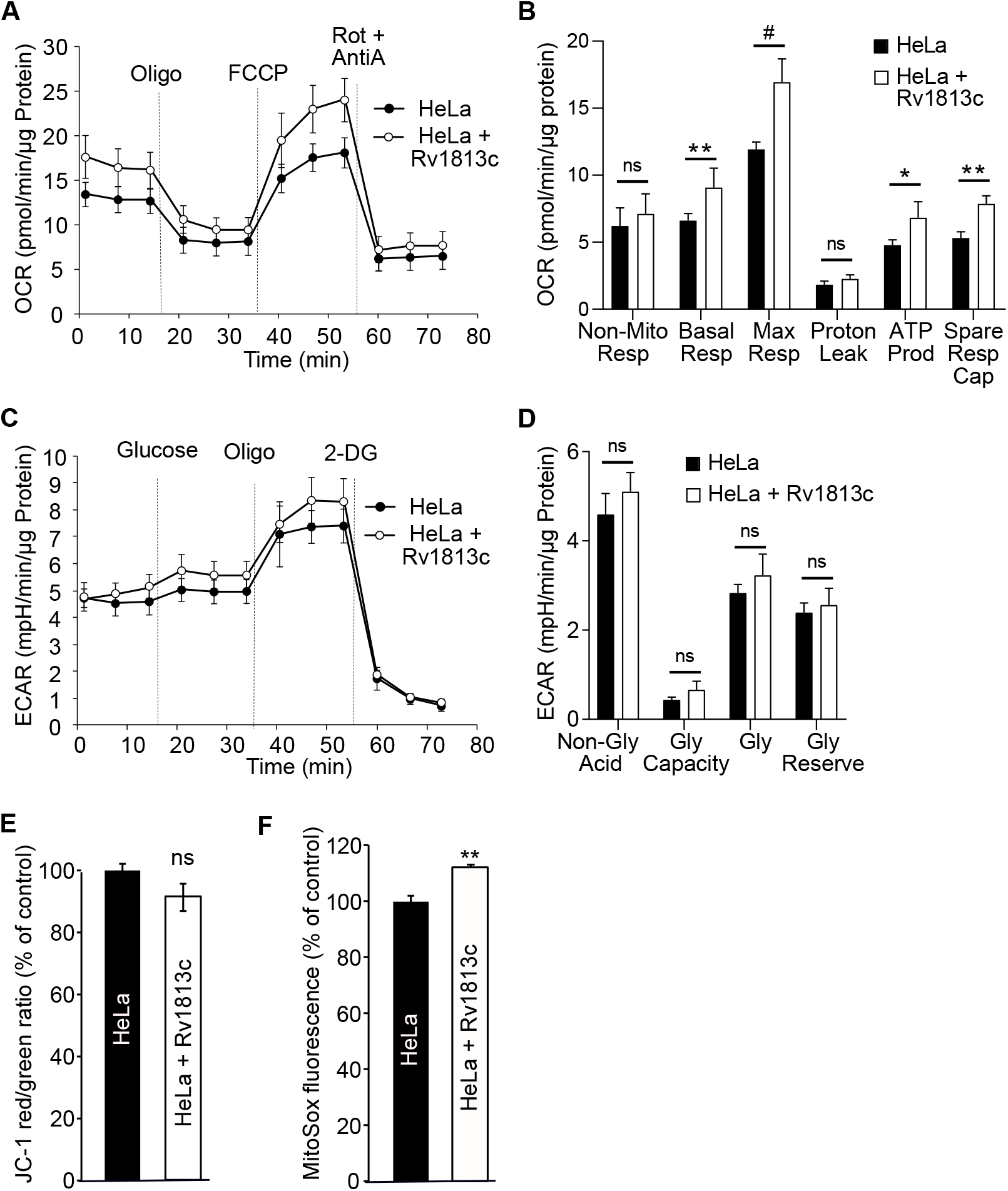
Functional consequences of Rv1813c mitochondrial localization. **(A,B)** Analysis of cell respiratory functions. HeLa cells were transiently transfected with pMACS-4-IRES-II Rv1813c or vector alone and enriched to >94% of expressing cells. Cell respiratory profiles (OCR) (A) and respiratory parameters (B) were obtained using an extracellular flux analyzer (Seahorse XF analyzer) and the mitochondrial respiration test. After reaching basal respiration, cells were subjected to 1 μM oligomycin to inhibit the ATP synthase and measure the mitochondrial ATP-linked OCR, followed by 1μM FCCP (cyanide-4-[trifluoromethoxy]phenylhydrazone) to uncouple mitochondrial respiration and maximize OCR, and finally 1 μM antimycin A (AntiA) and 100 nM rotenone (Rot) to inhibit complex III and I in the ETC respectively, and shut down respiration. In (B), analyzed respiratory parameters are non-mitochondrial respiration (Non-Mito Resp), basal respiration (Basal Resp), maximal respiration (Max Resp), proton leak, ATP production (ATP Prod) and spare respiratory capacity (Spare Resp Cap). **(C,D)** Analysis of glycolytic functions. Extracellular acidification (ECAR) profiles (C) and glycolytic parameters (D) of the same MACS enriched transfected cells were determined simultaneously to OCR analysis using the glycolysis stress test and the XF analyzer. After reaching non-glycolytic acidification, 10 mM glucose was added, followed by 1 μM oligomycin (Oligo) to inhibit the ATP synthase and induce maximal glycolysis. Finally, 100 mM 2-deoxyglucose (2-DG) was added to shut down glycolysis. This last injection resulted in a decreased ECAR confirming that the recorded ECAR was only due to glycolysis. In D, analyzed glycolitic parameters are non-glycolitic acidification (Non-Gly Acid), glycolytic capacity (Gly Capacity), glycolysis (Gly) and glycolytic reserve (Gly Reserve). Values are means ± sd. Student’s t test relative to HeLa cells #, p < 0.000001, ** p < 0.0005, * p < 0.005, ns non-significant. In **(E)** and **(F)**, HeLa cells were transiently transfected with pCI-myc-Rv1813c or empty vector as control and analyzed 48h later by flow cytometry **(E)** Flow cytometry analysis of the indicated HeLa cell lines stained with JC-1 to monitor mitochondrial membrane potential. JC-1 Red/Green ratio were calculated and expressed as the % of this ratio in HeLa cells. Values are means ± s.e.m. of three independent experiments. ns, not significantly different Student’s t-test. **(F)** Flow cytometry analysis of MitoSox stained HeLa cell lines. MitoSox fluorescence was expressed as the % of fluorescence in HeLa cells. Values are means ± s.e.m. of three independent experiments, ** p≤0.01 Student’s t-test.

### Oxidative stress-induced translocation of cytochrome c is delayed in Rv1813c expressing cells

We next assessed whether these mitochondrial alterations might alter the ability of Rv1813c expressing HeLa cells to cope with oxidative stress, a situation encountered during by *Mtb* infection. We choose to monitor the release of Cytochrome c (Cyt-c) from mitochondria into the cytosol in response to hydrogen peroxide, an early event in apoptotic cell death (Stridh et al., 1998). Hence, cells were incubated with hydrogen peroxide for three hours, and the localization of Cyt-c and Rv1813c was analyzed by confocal microscopy. As expected, Cyt-c showed a diffuse cytosolic staining in 21% of parental HeLa cells upon addition of 0.1mM hydrogen peroxide (**Fig. 6A, 6B**). Rv1813c release from mitochondria was also observed in cells overexpressing Rv1813c in response to hydrogen peroxide treatments (Fig. **6A**, **6C**). In contrast, Cyt-c release from mitochondria into the cytosol was reduced in Rv1813c expressing cells, with only 7.9% of cells displaying a cytosolic Cyt-c staining upon oxidative stress conditions (**Fig. 6A, B**). Note that cells with cytosolic Cyt-c always showed a strict concomitant Rv1813c cytosolic localization, thus we did not detect any cells harboring cytosolic Cyt-c and mitochondrial Rv1813c localizations together. Strikingly, Rv1813c release from mitochondria was more frequently observed than Cyt-c translocation leading to another cell population with Rv1813c in the cytosol but Cyt-c still in mitochondria (**Fig. 6D**). We conclude that the massive exit of Rv1813c from mitochondria in response to oxidative stress might delay the normal stress-induced Cyt-c translocation.

**Fig. 6:**
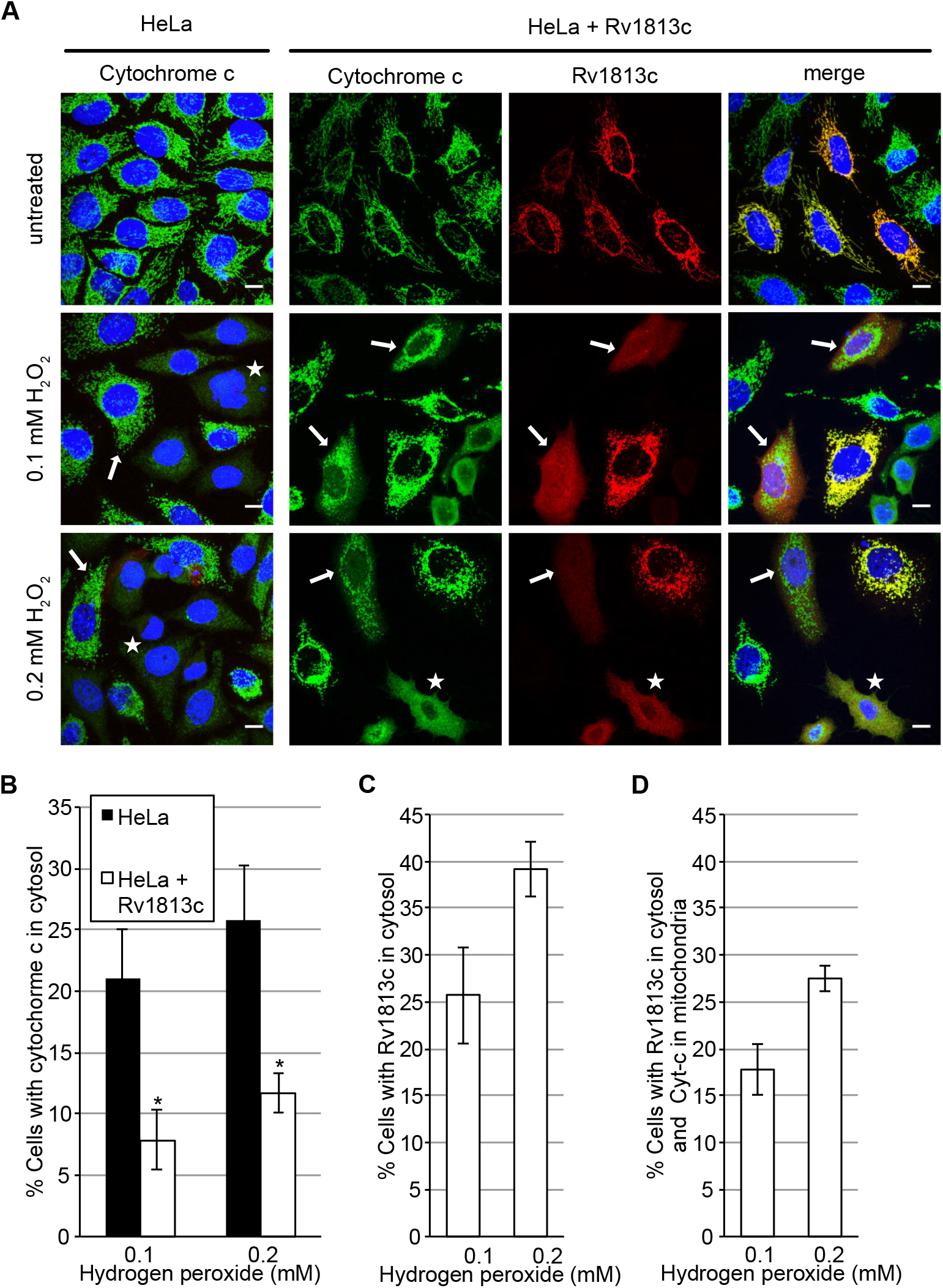
Analysis of Cyt-c and Rv1813c release from mitochondria upon oxidative stress. **(A)** Confocal microscopy analysis of HeLa cells transiently expressing myc-Rv1813. 48h post-transfection, cells were treated with 0.1 or 0.2 mM hydrogen peroxide for three hours, fixed, processed for immunofluorescence with anti-Cytochrome c (green) and anti-Rv1813c (red) antibodies, and observed by confocal microscopy. Nuclei were stained with Hoechst (blue). White arrows and white stars indicate cells with Cyt-c in mitochondria and cytosol respectively. Scale bar, 10 μm **(B, C, D)** Quantification of cells with Cyt-c in cytosol (B), with Rv1813c in cytosol (C) and Rv1813c in cytosol but Cyt-c in mitochondria (D) upon incubation with hydrogen peroxide for three hours. Values are means ± s.e.m. of three independent experiments, with 100 cells analyzed for each condition, * p<0.05 Student’s t-test.

## Discussion

Several intracellular pathogens (i.e. *Rickettsia, Legionella, Salmonella)* disrupt mitochondrial functions during infection mainly due to indirect effects (Spier et al., 2019)(Garaude, 2019) but more rarely by direct mitochondrial targeting of bacterial effectors having adverse functions (Hicks and Galán, 2013). For instance, the EspF effector from enteropathogenic *E. coli* is addressed to mitochondria via a mitochondrial import signal and promotes caspase-mediated apoptosis in intestinal epithelial cells (Hua et al., 2018). Furthermore, the MitF protein from *L. pneumophilia* was reported to alter mitochondria fission dynamics and promote a Warburg-like phenotype in macrophages (Escoll et al., 2017). Using bioinformatics screening and culture filtrate analysis, we have identified Rv1813c from *Mtb* as a secreted protein. The protein serves a role of vaccine adjuvant (Bertholet et al., 2008) and is highly immunogenic (Liang et al., 2019). The corresponding gene is non-essential for *Mtb* growth, however its deletion impairs *Mtb* virulence in a low dose murine model (Bretl et al., 2012). To our knowledge, no functional studies have been reported so far, thus the molecular basis of this attenuation is still unknown. Interestingly, Rv1813c expression is regulated by MprA and DosR proteins (Bretl et al., 2012). DosR is a transcriptional regulator induced by host intracellular stimuli, such as nitric oxide (NO), carbon monoxide (CO), and hypoxia (Bretl et al., 2012), while MrpA responds to environmental stress and residence within the macrophage (Haydel and Clark-Curtiss, 2004) (Pan et al., 2020) and is required during infection (Zahrt and Deretic, 2001). Accordingly, reference transcriptomes have revealed that Rv1813c is over-expressed (x2 and x4, 24h and 48h post-infection, respectively) in activated infected macrophages (Schnappinger et al., 2006), and in the BALB/c mouse model (x4, x5 then x6 respectively after 7, 14 and 21 days post-infection) (Talaat et al., 2004).

In this study, we established that Rv1813c belongs to a new protein family constitutively secreted by *Mtb* in culture medium and specifically addressed to mitochondria where it accumulates the IMS when ectopically expressed in host cells (**Fig. S1 and Fig. 2–4**). We solved the structure of Rv1813c that included a new protein fold with no similarity with any structures solved to date (**Fig. 1**). Interestingly, this small 9 kDa conserved folded domain was sufficient to specifically address the protein into mitochondria (**Fig. 2C**). Furthermore, we demonstrated that this localization subsequently enhances OXPHOS and inhibits cytochrome-c exit upon oxidative stress in Rv1813c-expressing cells. These Rv1813c-dependent phenotypes might be connected to important host defense mechanisms against *Mtb* infections and will have to be further addressed in *Mtb* infected cells.

Would the increased host cell mitochondrial ATP production induced by a *Mtb* effector protein bring any benefits to the bacteria for its intracellular replication or for avoiding normal host defense mechanisms? The activation of macrophages in response to *Mtb* infection is known to induce their polarization toward the toward an M1 profile (Shi et al., 2019). This important step is achieved by a metabolic reprogramming after NF-kB pathway activation either by pathogen-associated molecular patterns (PAMPs) or IFNg. NF-kB promotes the expression of the inducible nitric oxide synthase (iNOS) and subsequent nitric oxide (NO) release. Besides bactericidal activity, NO directly inactivates the electron transfer chain (ETC) proteins, triggering a complex series of events, mainly dependent on the production of reactive oxygen species (ROS) and change in the metabolites balance *(i.e.* NAD/NADP ratio). When the Krebs cycle is consequently blocked, citrate accumulates enhancing glycolysis and lipids biosynthesis. In addition, succinate also accumulates leading to HIF-1a (Hypoxia-inducible factor-1) stabilization, which completes this metabolic switch similarly to the Warburg effect observed in tumors (Shi et al., 2016). HIF-1a not only promotes the expression of enzymes involved in glycolytic ATP production, but also induces expression pattern leading to synthesis of important immune effectors, including inflammatory cytokines and chemokines under normoxic conditions (Wilson et al., 2019).

Very few studies have assessed the precise metabolic state of *Mtb* infected cells (Mohareer et al., 2020) and they have led to contradictory results. Recently, bioenergetic analyzes have been performed in infected macrophages. Hence, XF experiments and metabolites analysis have revealed a decrease of cell energetic flux through glycolysis and the TCA cycle in infected macrophages. Consequently, the total level of ATP produced in *Mtb* infected cells 5 and 24 hours post infection is reduced (Cumming et al., 2018). This observations confirm previous results indicating that the glycolytic flux was reduced in macrophages infected with virulent H37rv bacteria, a possible hallmark of a bacterial effector-induced incomplete or delayed metabolic shift (Simeone et al., 2012). On the contrary, other studies have suggested that maintaining host cell ATP production is beneficial for *Mtb* in order to avoid ROS production and apoptosis. For instance, a much higher ATP/ADP ration was observed in H37Rv-infected cells compared to cells infected with avirulent H37Ra (Jamwal et al., 2013)(Jamwal et al., 2016). In agreement, an elevated ATP/ADP ratio was also correlated to lower apoptosis rates observed in H37Rv-infected cells (Jamwal et al., 2013)(Mehrotra et al., 2014). Taken together, these data indicate that maintaining a high ATP production might be beneficial to delay a deleterious full metabolic shift and/or apoptosis of the host cell. Consistent with this hypothesis, secretion of Rv1813c could participate in maintaining a higher ATP production within mitochondria during *Mtb* infection.

Mitochondrial proton leak generated from the ETC is the major source of mitochondrial ROS. Excessive ROS amounts result in multiple effects including cytochrome-c translocation followed by caspase dependent apoptosis as well as inflammasome activation (Jamwal et al., 2013). Accordingly, the slight increase of ROS observed in resting Rv1813c-expressing cells (**Fig. 5F**) might be due ETC and/or ATP synthase boosted functions raising the ATP production in these cells. Interestingly, the artificial increase of ROS by exogenous hydrogen peroxide did not readily induce the release of Cyt-c from mitochondria in Rv1813c-expressing cells compared to parental HeLa cells. The release of Cytochrome c from mitochondria is an early event in apoptotic cell death and an early defense mechanism in infected *Mtb* macrophages (Li et al., 1997). Thus, this Rv1813c-based inhibition of this process might bring some advantages for *Mtb.* The molecular mechanism responsible for this inhibition will be further addressed.

In summary, this study provides some detailed suggestions for the possible regulatory functions of Rv1813c in the metabolic and apoptotic responses happening in *Mtb* infected macrophages.

**Detailed methods are provided in Supplementary Materials and Methods of this paper and include the following:**

◦ Purification of recombinant ^6^His-Rv1813c_28-143_ in *E. coli*
◦ Solution structure of Rv1813c_28-143_
◦ Antibodies
◦ Preparation of *Mycobacterium tuberculosis* culture
◦ Mycobacterial cell fractionation
◦ Protein Electrophoresis and Western Blot
◦ Cell culture and transfection conditions
◦ Mitochondria isolation and biochemical treatments
◦ Immunocytochemistry
◦ Flow cytometry analysis of JC-1, MitoSox and Annexin 5/PI stained cells
◦ MACS enrichment of CD4-Rv1813c transfected cells
◦ Extracellular flux analysis
◦ Transmission electron microscopy

## AUTHOR CONTRIBUTION

MM, AdV, PB, YMB, OP, TA, LH, JG, CG, GM, FL and MCG performed experiments; MM, GM, CG, ON, FL and MCG analyzed the data; FL and MCG conceived this study; All authors contributed to manuscript writing.

## AKNOWLEDGEMENTS

Flow cytometry and microscopy analyzes of uninfected cells were performed at the Montpellier RIO imaging facility of the University of Montpellier, member of the national infrastructure France-BioImaging, supported by the French National Research Agency (ANR-10-INBS-04, “Investments for the future”). The CBS acknowledges support from the French Infrastructure for Integrated Structural Biology (FRISBI) ANR-10-INSB-05-01. The following reagents were obtained through BEI Resources, NIAID, NIH: Monoclonal Anti-*M. tuberculosis* GlnA (Gene Rv2220), Clone IT-58 (CBA5) (produced *in vitro*), NR-44103; Polyclonal *Anti-Mycobacterium tuberculosis* FtsZ (Gene Rv2150c) (antiserum rabbit).

## DECLATION OF INTERESTS

The authors declare no competing interests.

## SUPPLEMENTARY DATA LEGENDS

**Fig. S1:**
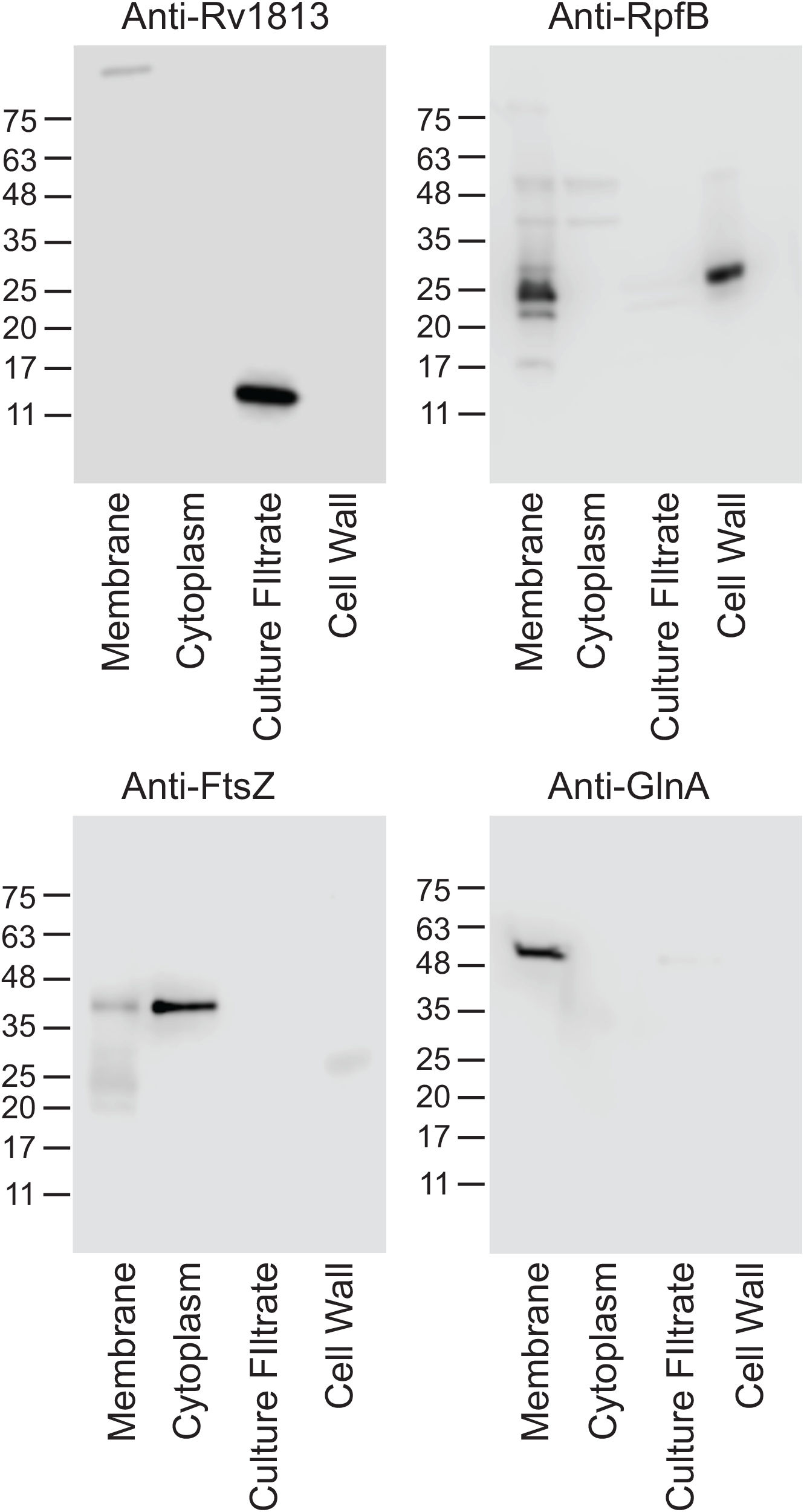
Rv1813 is detected in *M. tuberculosis* culture filtrate but not in cellular fractions. Lysate obtained from *M. tuberculosis* H37Rv strain grown in Sauton’s medium to logarithmic phase (OD_580_~0.7) were fractionated and probed with anti-Rv1813 polyclonal antibodies. Culture filtrates were obtained from the same culture. Anti-FtsZ (FtsZ is a cytoplasmic protein), anti-GlnA (GlnA is a membrane protein) and anti-RpfB (RpfB is a membrane and cell-wall anchored protein) antibodies were used to confirm the purity of mycobacterial fractions.

**Fig. S2:**
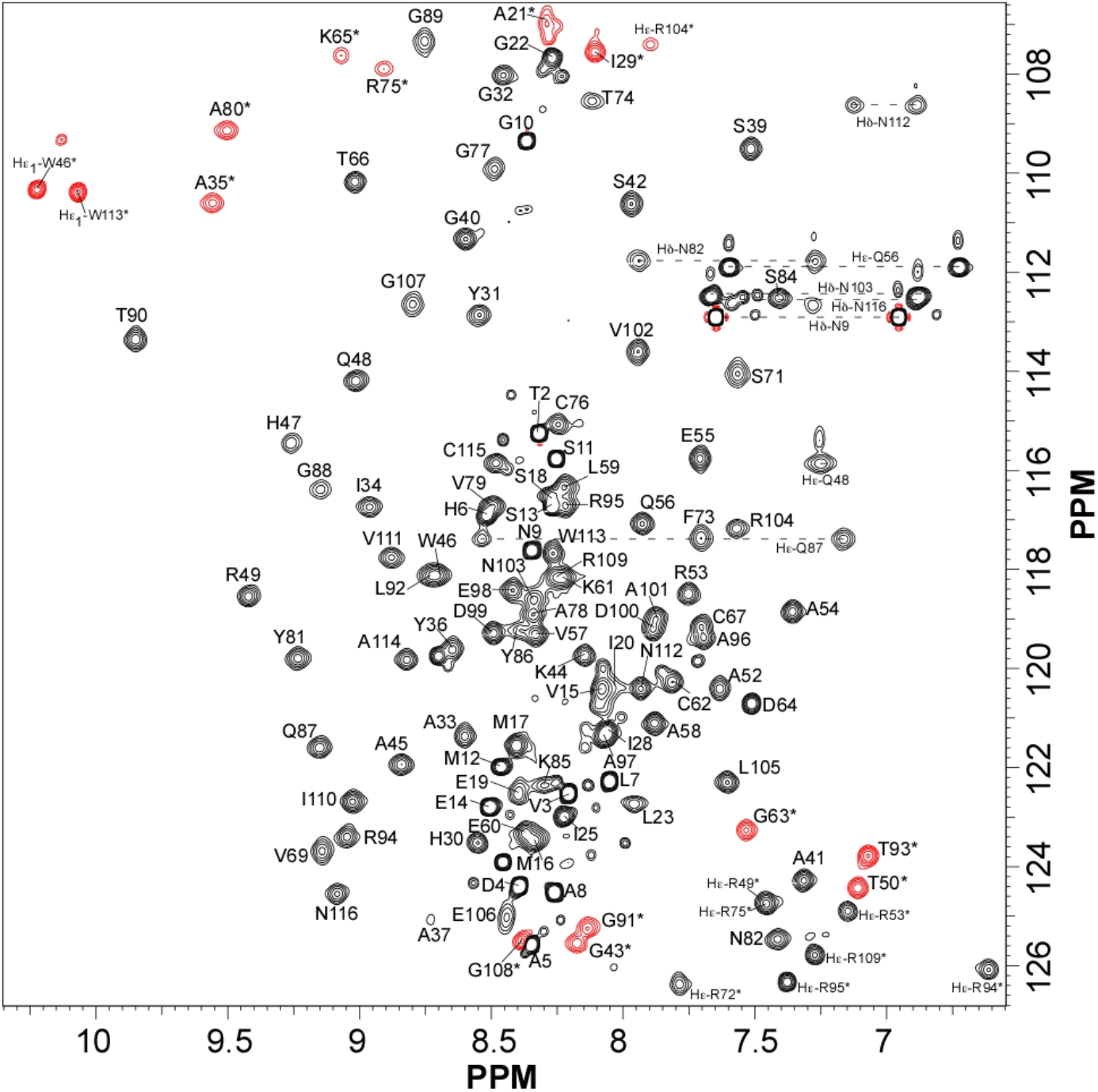
^1^H-^15^N HSQC spectrum of Rv1813c. This spectrum was obtained at 800 MHz, 20°C and pH 6,8 with 0.3 mM ^15^N-uniformly labeled sample. Cross peak assignments are indicated using the one-letter amino acid code and number following the full-length protein sequence numbering.

**Fig. S3:**
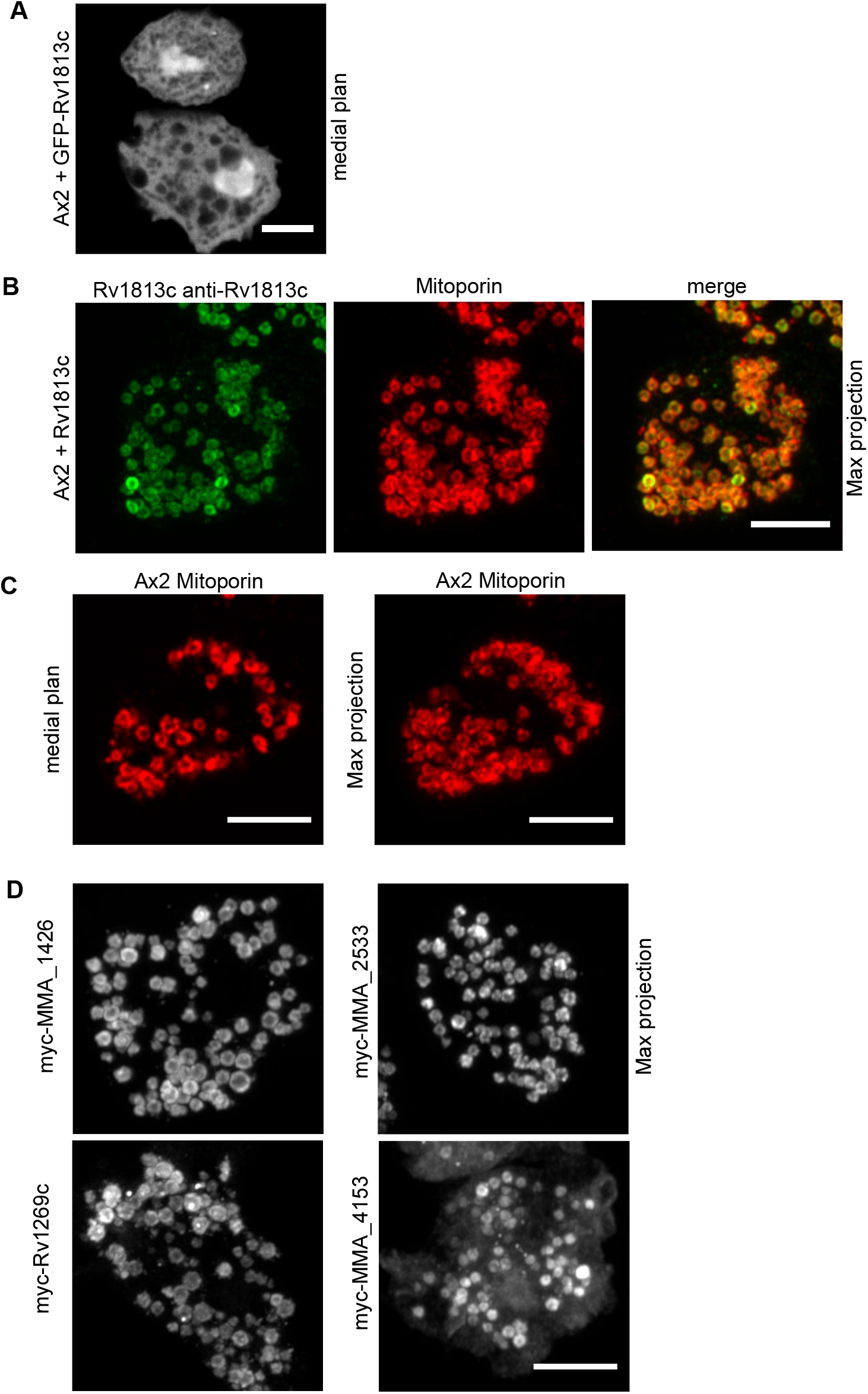
Confocal microscopy analysis of Rv1813c family members localization in *Dictyostelium.* *Dictyostelium* cells expressing the indicated constructs were fixed, processed for immunofluorescence, and analyzed by confocal microscopy (Airyscan). **(A)** Cellular localization of Nt-GFP-tagged Rv1813c. **(B)** Mitochondrial localization of untagged Rv1813c expressed in *Dictyostelium*. Rv1813c was labeled with a rabbit pAb anti-Rv1813c antibody. Rv1813c colocalizes with Mitoporin, a mitochondrial specific protein. **(C)** Mitoporin localization in untransfected parental Ax2 cells. Cells were labeled with a mouse mAb to mitoporin revealing characteristic ring shaped structures. A maximum projection of Z confocal sections is shown on the right panel. **(D)** Localization of different Rv1813c family members of *M. tuberculosis* and *M. marinum* expressed in *Dictyostelium and* revealed by anti-myc labelling. White arrows indicate so mitochondria with affected shapes. Bar, 5 μm.

**Fig. S4:**
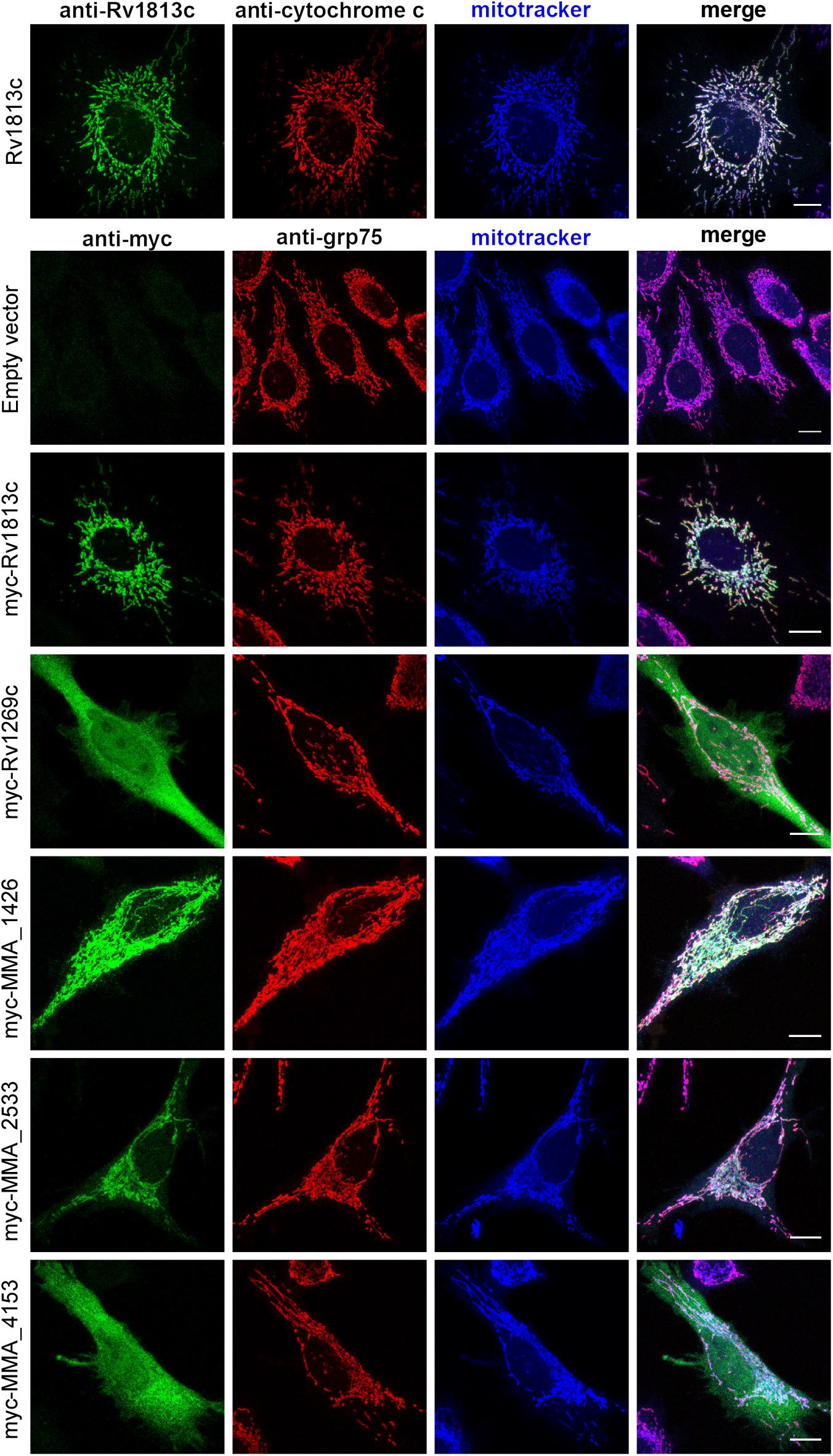
Rv1813c family localization in HeLa cells. HeLa cells expressing the indicated constructs were fixed, processed for immunofluorescence, and analyzed by confocal microscopy (Airyscan). Cells were colabeled either with rabbit polyclonal anti-Rv1813c, mouse mAb anti-Cytochrome c, and mitotracker deep red (upper panel) or anti-myc, rabbit anti-grp75 (mitochondria marker) and mitotracker deep red (lower panels). Bar, 10 μm.

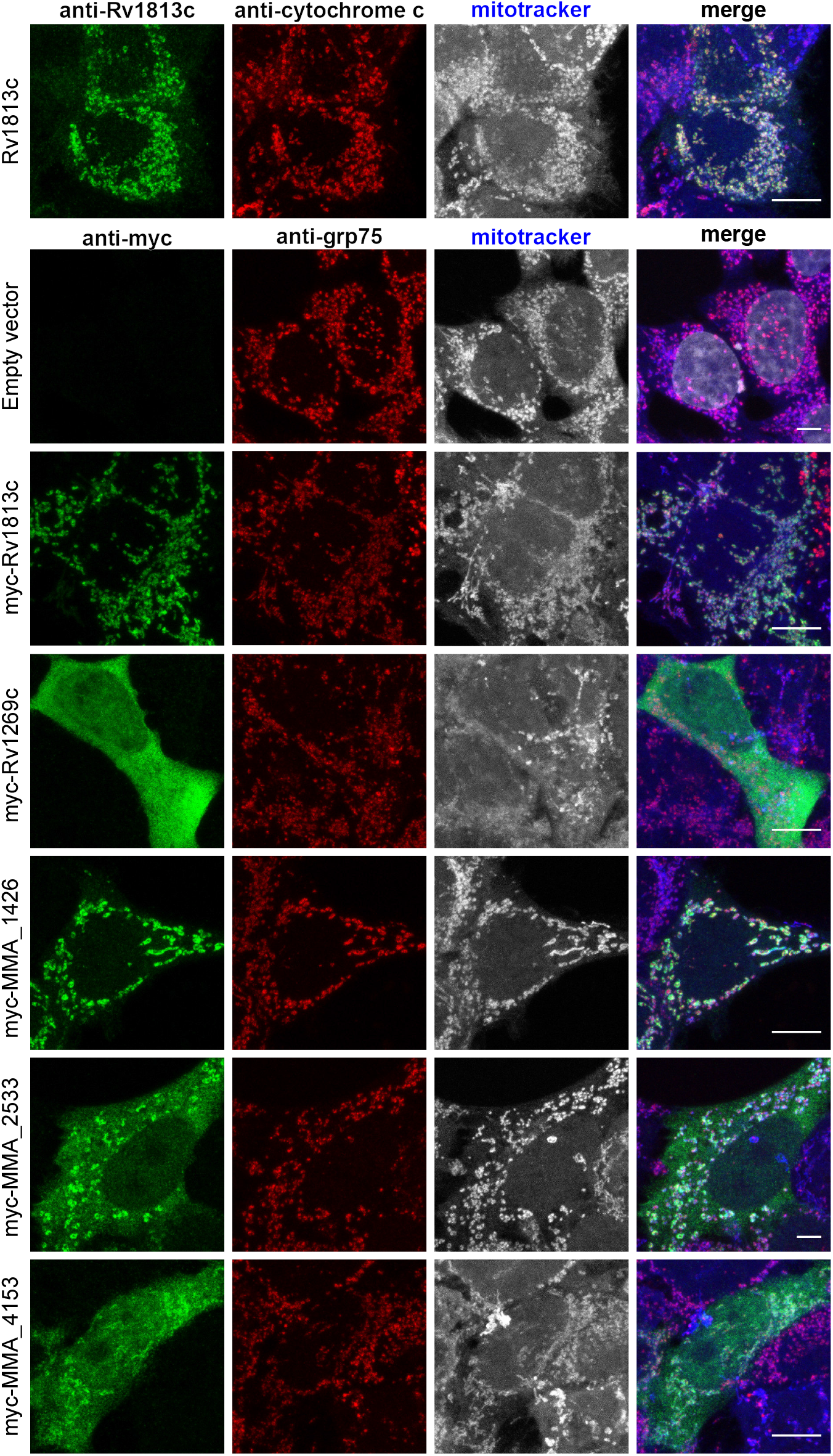

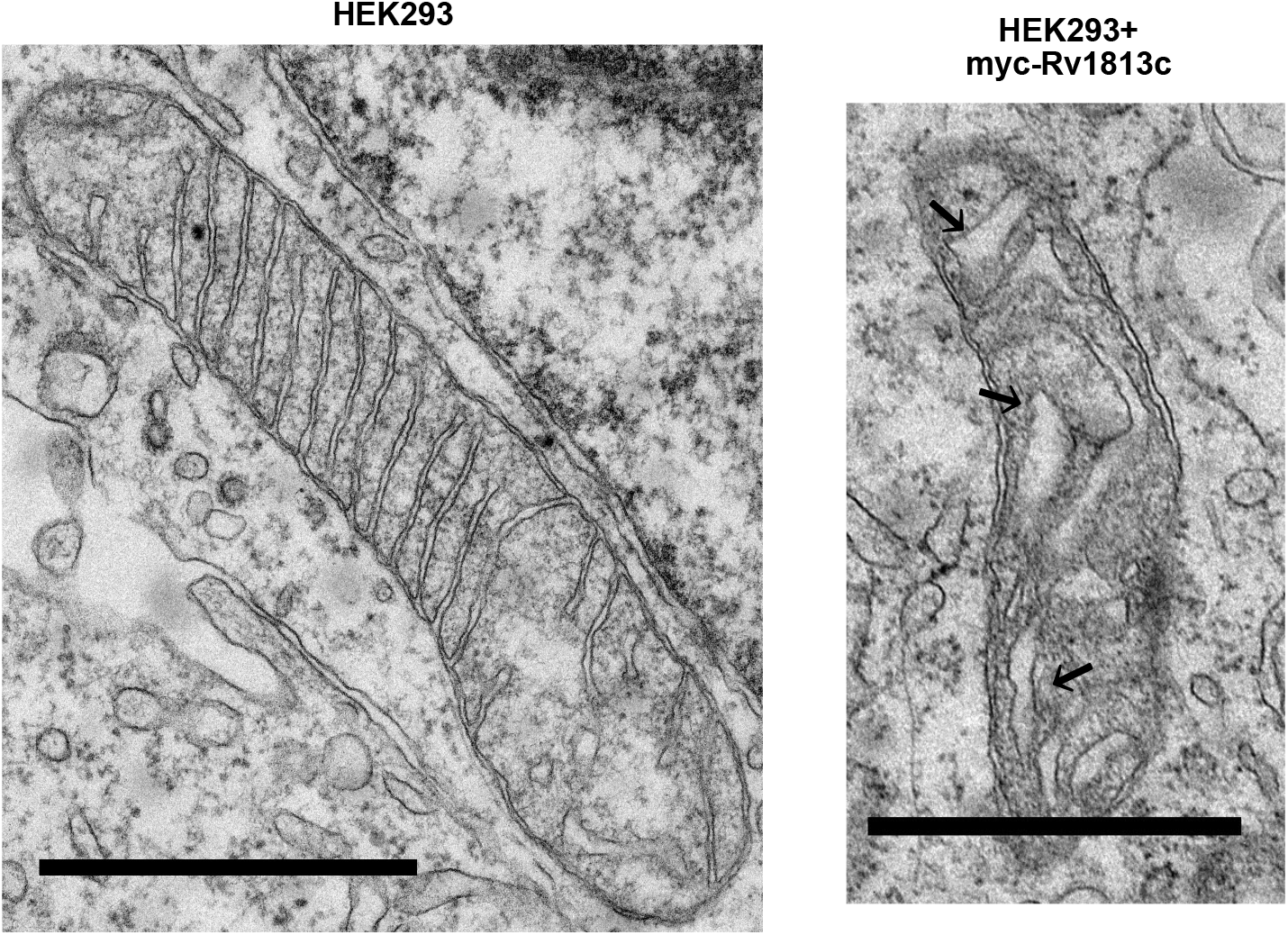

**SUPPLEMENTARY TABLE 1.**
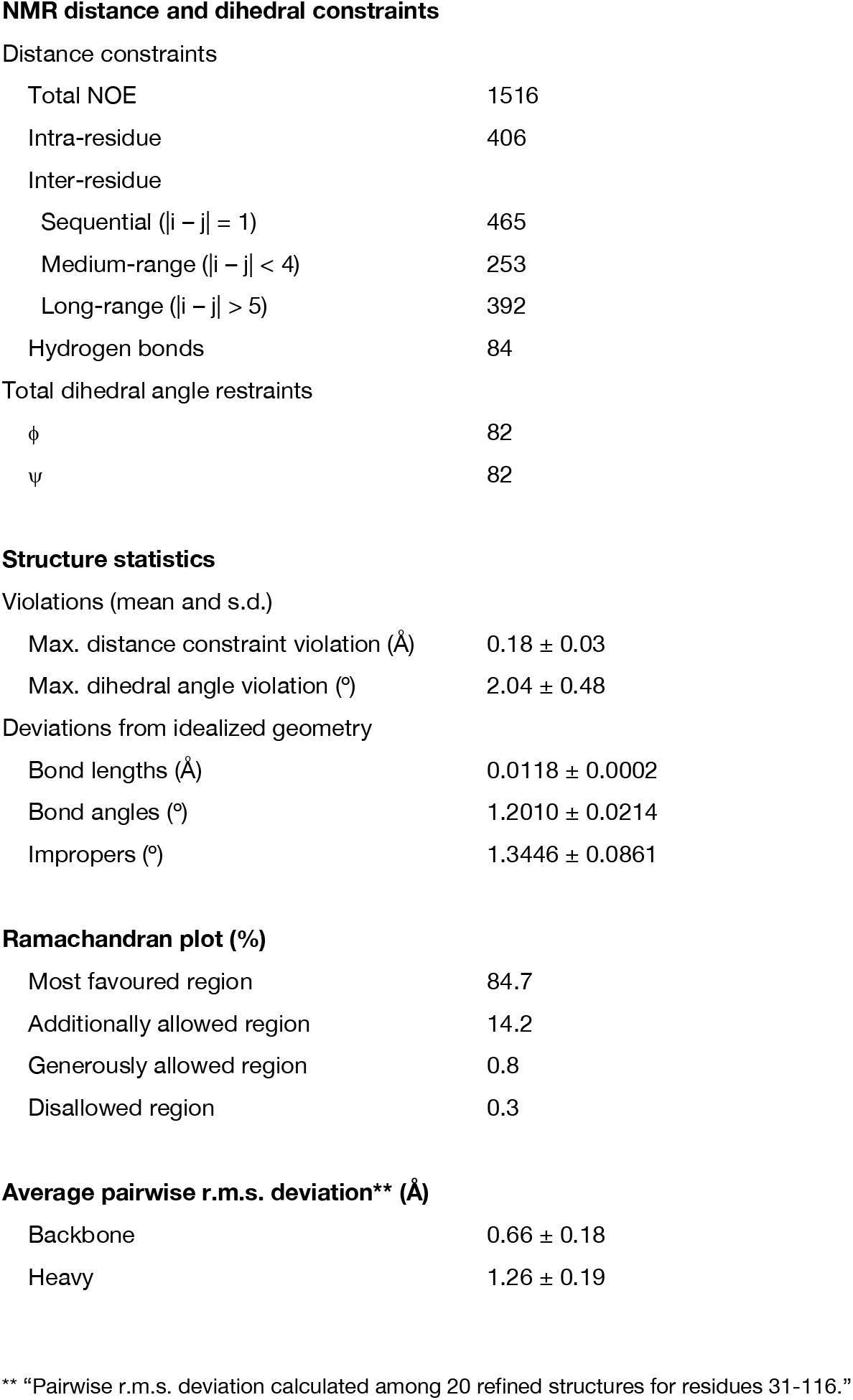
NMR and refinement statistics for RV1813 protein structures.

## Supplementary Materials and methods

### Purification of recombinant ^6^His-Rv1813c_28-143_ in *E. coli*

*E. coli* BL21(DE3) strains containing pET::*rv1813*_28-143_ vector were used to inoculate 1 L of LB medium supplemented with ampicillin (100 μg/ml) and resulting cultures were incubated at 37 °C with shaking until A_600_ reached ~0.5. Then, 1 mM final of isopropyl 1-thio-β-d-galactopyranoside was added and growth was continued for 3 hr at 37 °C. The cells were harvested by centrifugation and the resulting cell pellet was resuspended in buffer A (50 mM Tris-HCl pH 8.5, 150 mM NaCl, 2mM DTT). Cells were then lysed by sonication and cell debris and insoluble materials were separated by centrifugation. The pellet was then resuspended in buffer B (Buffer A + 8M Urea). After centrifugation the supernatant was loaded into a Hitrap™ IMAC HP column (Amersham biosciences), equilibrated in buffer B and 4 % of buffer C (buffer B supplemented with 300 mM of imidazole). The column was washed with successive applications of buffer B (approximately 30 ml in total) to remove all the impurities and then buffer C was increased over 20 ml to 100%. Fractions containing the Rv1813c proteins were identified by SDS-PAGE, then pooled and concentrated using a 5 K cut-off concentrator to a 2mg/ml concentration. The protein was dialysed against buffer A over-night at 4°C. The refolded protein was very unstable until removal of the 6His tag using 3C protease (4h digestion at 4°C). The protein was then loaded to a Superdex 75 26/60 (Amersham biosciences) size exclusion column, equilibrated in buffer 20 mM Na-Phosphate pH 6.2, 150 mM NaCl. Again, fractions containing the protein were identified by SDS-PAGE, then pooled and stored at −20°C until required. This protocol was carried out for all the non-labelled constructs of Rv1813c as well as for ^15^N and ^15^N -^13^C labelled constructs, except that the cultures were grown in a minimum media containing ^15^NH_4_Cl and ^15^NH_4_Cl/^13^C6-glucose as the sole nitrogen and carbon sources.

### Solution structure of Rv1813c (residue 28-143)

All NMR experiments were generally carried out at 25°C on Bruker Avance III 700 (^1^H-^15^N double resonance experiments) or Avance III 500 (^1^H-^13^C-^15^N triple-resonance experiments) spectrometer equipped with 5 mm z-gradient TCI cryoprobe, using the standard pulse sequences. NMR samples consist of approximately 0.9 mM ^15^N- or ^15^N,^13^C-labeled protein dissolved in 25 mM NaCitrate, 150 mM NaCl (pH 5.6) with 10% D_2_O for the lock. ^1^H chemical shifts were directly referenced to the methyl resonance of DSS, while ^13^C and ^15^N chemical shifts were referenced indirectly to the absolute ^15^N/^1^H or ^13^C/^1^H frequency ratios. All NMR spectra were processed and analyzed with GIFA. Backbone and Cβ resonance assignments were made using standard HNCA, HNCACB, CBCA(CO)NH, HNCO, and HN(CA)CO experiments performed on the ^15^N,^13^C-labeled Rv1813c_28-143_ sample. NOE cross-peaks identified on 3D [^1^H, ^15^N] NOESY-HSQC (mixing time 120 ms) were assigned through automated NMR structure calculations with CYANA 2.1, whereas NOE on 3D [^1^H,^13^C] NOESY-HSQC were assigned manually. Backbone φ and ψ torsion angle constraints were obtained from a database search procedure on the basis of backbone (^15^N, H_N_, ^13^C’, ^13^Cα, Hα, ^13^Cβ) chemical shifts using the program TALOS+ (Shen et al., 2009). Hydrogen bond restraints were derived using standard criteria on the basis of the amide ^1^H / ^2^H exchange experiments and NOE data. When identified, the hydrogen bond was enforced using the following restraints: ranges of 1.8–2.0 Å for d(N-H,O), and 2.7-3.0 Å for d(N,O). The final list of restraints, from which values redundant with the covalent geometry has been eliminated. The 30 best structures (based on the final target penalty function values) were minimized with CNS 1.2 according the RECOORD procedure (Nederveen et al., 2005) and analyzed with PROCHECK (Laskowski et al., 1993). The rmsds were calculated with MOLMOL (Koradi et al., 1996). All statistics are given in Table 1. The chemical shift table was deposited in the BMRB databank (accession number 7NHZ).

### Antibodies

The following primary antibodies were used in this study: mouse anti-Myc (Invitrogen, #13-2500, 1:200 for immunofluorescence, 1:500 for immunoblot), mouse anti-cytochrome c (clone 6H2.B4, BD PharMingen, 1:500 for immunofluorescence), mouse *anti-Dictyostelium* Mitoporin (70-100-1; 1:2000 for immunofluorescence and immunoblot) (Troll *et al.,* 1992), rabbit anti-Rv1813c raised using recombinant Rv1813c (ProteoGenix SAS, Schiltigheim, France) (1:2000 for immunofluorescence, 1:5000 for immunoblot), rabbit anti-Grp75 (D13H4, XP #3593, Cell Signalling, 1:100 for immunofluorescence), rabbit anti-EHD (Dias *et al.,* 2012; 1:4000 for immunoblot). Secondary antibodies used for immunoblotting were horseradish peroxidase (HRP)-conjugated donkey anti-mouse IgG (H+L) (#715-035-151) and HRP-conjugated donkey anti-rabbit IgG (H+L) (#715-035-152) (Jackson ImmunoResearch). MCCC1 (mitochondrial matrix) was revealed by staining with HRP-conjugated streptavidin as previously described (Davidson et al., 2013). Secondary antibodies used for immunofluorescence were Alexa-Fluor-568-conjugated goat anti-mouse IgG (H+L) (#A11031), Alexa-Fluor-594-conjugated donkey anti-rabbit IgG (H+L) (#A21207), Alexa-Fluor-488-conjugated goat anti-rabbit IgG (H+L) (#A11029) and Alexa-Fluor-488-conjugated donkey anti-rabbit IgG (H+L) (#A21206) (ThermoFisher Scientific, Illkirsh, France). All secondary antibodies were used at 1:500 for immunofluorescence. Prolong Golf Antifade and Hoechst 33342 (#62249) were purchased from Molecular Probes (ThermoFisher Scientific, Illkirsh, France).

### Preparation of *Mycobacterium tuberculosis* culture

*M. tuberculosis* was grown in Middlebrook 7H9 liquid medium supplemented with 10% (v/v) Albumin-Dextrose Complex (ADC), 0.2% (v/v) glycerol and 0.1% Tween 80 (w/v), at 37°C in a roller incubator. Bacterial growth was followed by measurement of absorbance at 580 nm, using a spectrophotometer, or by colony-forming unit (CFU) counting on 7H10 agar.

### Mycobacterial cell fractionation

Mycobacteria cell fractionation was done as described else were (O.Turapov, Cell Report, 2018). Briefly, cells were lysed in a buffer that contained 20 mM TrisHCl, pH 8.0, 150 mM NaCl, 20 mM KCl, 10 mM MgCl2. Bacterial culture was homogenized with a Minilys homogenizer (Bertin Instruments) using glass beads. A cocktail of proteinase/phosphatase inhibitors (Roche, UK) were used in all buffers. Lysates were centrifuged for 1 hour at 27,000 x g, the pellets were washed in a carbonate buffer (pH 11) and used as a cell wall material. The supernatant was centrifuged again for 4 hours at 100,000 x g. The supernatants from this step was used as cytoplasmic fraction and the pellets (membrane fractions) were washed once in carbonate buffer, pH 11 and twice in TBS buffer. Proteins from cellular fractions were separated on SDS-PAGE. The purity of fractions was confirmed by the detection of diagnostic proteins as described below.

### Protein Electrophoresis and Western Blot

Proteins were separated on 4%–20% gradient SERVA gels and transferred onto a nitrocellulose membrane using a Trans-Blot^®^ Turbo Transfer System (Bio-Rad) according to the manufacturer’s instruction. SignalFire Elite ECL Reagent (Cell Signalling, UK) were used to visualize proteins on C-DiGit Chemiluminescent Blot Scanner (LI-COR Biosciences), according to the manufacturer’s instructions. All the secondary antibody were from Cell Signalling, UK. Diagnostic proteins were used for all the cellular fractions: GlnA (membrane protein), GarA (secreted and cytoplasmic protein), RpfB (membrane and cell wall protein) and FtsZ (cytoplasmic protein).

### Cell culture and transfection conditions

*D. discoideum* strain Ax2 was grown at 22°C in HL5c medium supplemented with 18 g/L Maltose (Formedium, Norfolk, United Kingdom). For ectopic expression in *Dictyostelium,* Rv1813c family coding sequences with *Dictyostelium* optimized codons (IDT, Integrated DNA Technologies, Inc., Coralville, Iowa 5224, USA) were cloned into pDXA-3C-myc (Manstein *et al.,* 1995). Plasmids were linearized by ScaI and transfected by electroporation as described (Cornillon *et al.,* 2000). Clones were selected in 5μg/mL G418.

HeLa (ATCC CRM-CCL-2) and HEK-293T (ATCC CRL-3216) cells were maintained in DMEM, high glucose (Dulbecco’s Modified Eagle Medium) containing 5% and 10% heat-inactivated foetal bovine serum, respectively, and supplemented with GlutaMAX™ (Gibco Life Technologies), penicillin (100 units/mL), and streptomycin (100 μg/mL). Transfections of HeLa cells were performed using JetPEI™ transfection reagent (PolyPlus-Transfection, Ozyme, Saint Quentin, France), according to the manufacturer. Cells plated one day before transfection were incubated with JetPEI™ -DNA complexes (N/P=5), and after 5h the medium was changed. All assays were performed 48h post-transfection.

For confocal microscopy analysis, HeLa cells were seeded on glass coverslips coated with 0.001% poly-L-Lysine (# P4707, Sigma). For localization, Rv1813c family coding sequences with human optimized codons were cloned into the mammalian expression vector pCI (a kind gift of Dr. Solange Desagher, IGMM, Montpellier, France). Cells on glass coverslips were transfected in a 24-well culture plate and analyzed 48h later. For mitochondrial membrane potential, mitochondrial ROS and oxidative stress studies, cells were transfected on 6-well culture plates. After 24h, resuspended cells were pooled and plated either on glass coverslips for confocal microscopy or on 6-well culture plates at a density of 2-3.10^5^ cells/well for FACS analysis. For extracellular flux analysis, HeLa cells seeded into five 100-mm tissue culture dishes were transfected with Rv1813c DNA cloned into pMACS 4-IRESII vector (Miltenyi Biotec, France), allowing Rv1813c co-expression with a truncated CD4 surface marker. After 24h, EDTA resuspended cells were pooled and CD4 positive cells selected through magnetic cell sorting (MACS) as described below.

### Mitochondria isolation and biochemical treatments

Mitochondria were isolated as described (Aubry *and* Klein, 2006). Briefly *Dictyostelium* cells were washed in ice-cold buffer A (20 mM HEPES pH7, 1 mM EDTA, 250 mM Sucrose, proteinase inhibitors), resuspended at a cell density of 3×10^8^ cells/mL, and broken with a ball bearing homogenizer (8.02 mm bore, 8.002 mm ball; 20 strokes). Unbroken cells were removed by low speed centrifugation (5 min, 1500 g). The supernatant was next centrifuged for 15 min at 16,000 g. The pellet was resuspended in buffer A and the centrifugation repeated to yield the enriched mitochondria fraction. For further subcellular fractionation, this fraction was further centrifuged at 100,000 g for 1h. Triton X-114 phase fractionation was performed as described (Bordier, 1981). Briefly, mitochondria were incubated for 20 min at 4°C in 10 mM Tris-HCl pH7.4, 150 mM NaCl and 1% Triton X-114. Samples were loaded on a 6% sucrose cushion, incubated at 30°C for 3 min for condensation, and centrifuged at 300 g for 3 min at room temperature. Supernatants were adjusted to 1% Triton X-114 and the procedure repeated. Detergent and aqueous phases were analyzed by western blotting.

For Carbonate extraction of integral membrane proteins, mitochondria were incubated for 30 min at 4°C in 0.1 M Na_2_CO_3_ pH11.5 and centrifuged for 30 min at 100,000 g as previously described (Fujiki et al., 1982).Pellets were resuspended in buffer A. Proteins in resuspended pellets and supernatants were precipitated with 15% TCA and resuspended in SDS page loading buffer. Integral membrane proteins were recovered in the pellet, while soluble and peripheral proteins were present in the supernatant. For high salt washes, intact mitochondria were incubated in 10 mM Tris-HCl pH7.3, 250 mM Sucrose, 200 mM KCl and incubated for 30 min at 4°C. Mitochondria were then centrifuged for 10 min at 16,000 g. Pellets and supernatants were treated as above. For proteinase K digestions of mitochondrial peripheral membrane proteins, mitochondria in 20 mM HEPES pH7, 250 mM Sucrose, 100 mM KCl, 2 mM MgCl_2_, 1mM KH_2_PO_4_ were incubated with 100 μg/mL proteinase K for 30 min at 4°C ± 1% Triton X100. Samples were then treated with TCA for protein precipitation. To break selectively mitochondrial outer membranes, mitochondria were resuspended in hypotonic buffer (2 mM HEPES pH7, 5 mM KCL, proteinase inhibitors) for 30 min at room temperature. After centrifugation at 16,000 g for 10 min, pellets and supernatants were treated with TCA as above.

### Immunocytochemistry

*Dictyostelium* cells were applied on glass coverslips for 3h, and then fixed with 4% paraformaldehyde for 30 min, washed and permeabilized for 2 min in −20°C methanol. Cells were incubated with the indicated antibodies for 1h, washed, and then stained with appropriate fluorescent secondary antibodies for 30 min. After three washes, coverslips were mounted in Mowiol. Mammalian cells were cultured on glass coverslips and fixed with 4% paraformaldehyde in phosphate-buffered saline (PBS) for 20 min. Cells were washed in Tris-buffered saline (TS; 25mM Tris pH7.4, 150mM NaCl) for 10 min. After permeabilization with 0.2% Triton X-100 in TS for 4 min, non-specific binding was blocked with 0.2% gelatin from cold water fish skin (Sigma-Aldrich, France) in TS for 30 min. Cells were incubated with primary antibodies in blocking buffer for 1h and were then washed 3 times with 0.008% TritonX-100 in TS for 10 minutes. Cells were incubated for 30 minutes with Alexa-Fluor-labelled secondary antibodies in blocking buffer. After rinsing in washing buffer, cell nuclei were stained with 1 μg/ml Hoechst in TS for 5 minutes. Finally, coverslips were mounted with Prolong Gold Antifade (#P36934 Thermo Fisher Scientific). Slides were examined under a Leica TCS SPE confocal microscope equipped with a 40X/1.15 or 63X/1.33 ACS APO oil-immersion objective or a Zeiss LSM880 AiryScan confocal microscope equipped with a 40X/1.4 or 63x/1.4 Oil Planapochromat DIC objective. Fluorescence images were adjusted for brightness, contrast and colour balance by using the ImageJ software.

### Flow cytometry analysis of JC-1 and MitoSox stained cells

For MitoSox red staining of HeLa cells, 2.5×10^5^ cells resuspended in CPBS buffer (PBS, 2.67 mM KCl, 0.5 mM MgCl_2_, 0.7 mM CaCl_2_ and 0.1% glucose) were incubated in 5 μM MitoSox red. After 20 min at 37°C with shaking, cells were washed twice in CPBS buffer before FACS analysis. JC-1 staining of HeLa cells was made according to the manufacturer recommendations. Briefly, cells cultured in 6-well culture plates (2.5×10^5^/well) were incubated at 37°C in culture medium supplemented with 2 μM JC1. After 30min, cells were washed, resuspended in PBS, and directly analyzed by flow cytometry. As positive control of JC-1 staining, 5 μM carbonyl cyanide m-chlorophenyl hydrazone (CCCP) was added to cells during JC-1 cell incubation.

### MACS enrichment of CD4-Rv1813c transfected cells

MACS enrichment of transfected cells was done with MACSelect Transfected Cell Selection kit from Miltenyi Biotec, according to the supplier. Briefly, HeLa cells were transfected with empty pMACS4-IRESII (as control) or pMACS4-IRESII-Rv1813c plasmids allowing expression of truncated CD4 cell surface marker alone or in combination with Rv1813c respectively. After 24h, ~10^7^ cells were washed, dissociated in ice cold PBS containing 5 mM EDTA, centrifuged at 200 g for 10 minutes at 4°C, and resuspended in 320 μl ice-cold de-gassed PBS supplemented with 0.5% bovine serum albumin and 5 mM EDTA (PBE). Magnetic labelling of the transfected cells was achieved by incubating cells with 80 μl of anti-CD4 coupled MACSelect MicroBeads on ice for 15 minutes. Volume was adjusted to 2 ml with PBE and cells were subjected to magnetic separation using LS column (Miltenyi Biotec) and MACS separator. After three washes with 3 ml of PBE, cells were flushed out with 5 ml of PBE. To increase the purity of the magnetically labelled fraction, magnetic separation was repeated once on a second LS column. After the final wash, cells were flushed out with 5ml of cell culture medium, counted and seeded at a density of 1.85×10^4^ cells/well on XF96 cell culture microplates (Seahorse, Agilent Technologies, France) previously coated with 0.1mg/ml poly-D-Lysine (#P7280, SIGMA) or on glass coverslips to evaluate the level of MACS enrichment of transfected cells by immunofluorescence. Cells were incubated at 37°C and analyzed 24h later using the Seahorse XF96 extracellular flux analyzer or by confocal microscopy.

### Extracellular flux analysis

Cells plated the day before on XF96 cell culture microplates were washed with pre-warmed cell culture medium 5h before analysis to eliminate dead cells. Extracellular Flux analysis was performed using Seahorse XF Extracellular Flux analyzer, allowing simultaneous measurement of oxygen consumption rate (OCR) and extracellular acidification rate (ECAR). Mitochondrial respiration and glycolytic function of the cells were measured using Cell Mito Stress Test Kit (#103015-100) and Cell Glycolysis Stress Test Kit (#103020-100), respectively (Agilent Technologies, France). Cells were incubated in Seahorse XF DMEM pH7.4 (#103575-100, Agilent) supplemented with 1 mM sodium pyruvate, 2 mM glutamine and with 10 mM glucose (Cell Mito Stress Test Kit) or without glucose (Cell Glycolysis Stress Test Kit) in a 37°C incubator without CO_2_ for 1h prior to the assay. After calibration and three initial measurements at baseline, different perturbing chemicals corresponding to each kit were sequentially injected, and three successive measurements were taken after each injection.

### Transmission electron microscopy

MACS enriched cells on glass coverslips were successively fixed with 2.5% gluteraldehyde in 0.1 M cacodylate buffer pH 7.4, washed with cacodylate buffer, post-fixed in 1% osmium tetroxide in cacodylate buffer, washed with distilled water, and finally incubated in 1% uranyl acetate. Dehydration was performed through acetonitrile series. Samples were impregnated first in epon 118: acetonitrile 50:50, and twice in 100% epon. After overnight polymerization at 60°C, coverslips were detached by thermal shock with liquid nitrogen. Polymerization was then prolonged for 48h at 60°C. Ultrathin sections of 70 nm were cut with a Leica UC7 ultramicrotome (Leica microsystems), counterstained with lead citrate and uranyl acetate prepared in ethanol. Sections were observed in a Jeol 1200 EXII transmission electron microscope. All chemicals were from Electron Microscopy Sciences (USA) and solvents were from Sigma. Images were processed using the Fiji software.

